# A peptide-level multiple imputation strategy accounting for the different natures of missing values in proteomics data

**DOI:** 10.1101/2020.05.29.122770

**Authors:** Q. Giai Gianetto, S. Wieczorek, Y. Couté, T. Burger

## Abstract

**Motivation:** Quantitative mass spectrometry-based proteomics data are characterized by high rates of missing values, which may be of two kinds: missing completely-at-random (MCAR) and missing not-at-random (MNAR). Despite numerous imputation methods available in the literature, none account for this duality, for it would require to diagnose the missingness mechanism behind each missing value.

**Results:** A multiple imputation strategy is proposed by combining MCAR-devoted and MNAR-devoted imputation algorithms. First, we propose an estimator for the proportion of MCAR values and show it is asymptotically unbiased under assumptions adapted to label-free proteomics data. This allows us to estimate the number of MCAR values in each sample and to take into account the nature of missing values through an original multiple imputation method. We evaluate this approach on simulated data and shows it outperforms traditionally used imputation algorithms.

**Availability:** The proposed methods are implemented in the R package imp4p (available on the CRAN Giai Gianetto (2020)), which is itself accessible through Prostar software.

## 1 Introduction

A widely used method for identifying and quantifying broad amounts of proteins in biological samples is based on the label free mass spectrometry (MS) analysis of their constituting *peptides* (protein fragments obtained by enzymatic digestion). This experimental pipeline is particularly suitable for discovery proteomics for its high proteome coverage and throughput when compared to pipelines either relying on isotopic labeling or analyzing intact proteins (Nesvizhskii and Aebersold, 2005). Unfortunately, it comes at the price of lower data quality. Notably, the resulting peptide-level datasets are impacted by high rates of missing values (usually between 20% and 50%, see Webb-Robertson *et al.* (2015)), which are known to be of two kinds (Karpievitch *et al.* (2009)): Missing Not-At-Random (MNAR), mostly coming from the various phenomena that impact the lower detection limit of the mass spectrometer, and Missing Completely-At-Random (MCAR), resulting from the pipeline intrinsic non-exhaustiveness. This missing value concern is ubiquitous to all biomolecule analyses using label free MS, as in metabolomics for instance (Wei *et al.*, 2018). Therefore, although this article focuses on peptide datasets, the proposed methodology can be directly applied to other types of MS-based omics data.

Despite the many existing imputation methods (see Webb-Robertson *et al.* (2015) survey), there is still no consensus on how to proceed with missing values in label free MS. No method accounts for both MNARs and MCARs, so that the practitioner has to make an arbitrary choice on the missing value mechanism when imputing data. Yet, applying imputation strategies that treat all missing values in the same way regardless of their nature can lead to distorting reality and thus compromising the veracity of any biological conclusions that may result (Lazar *et al.* (2016)). Although some works proposed to account for different types of missing values (Luo *et al.*, 2009; Taylor *et al.*, 2013; Ryu *et al.*, 2014; Chen *et al.*, 2014; O’Brien *et al.*, 2018), they rely on single models describing the joint impact of the various missingness mechanisms without relying on imputation. In these studies, the model is directly used to find differentially abundant proteins and to infer new biological knowledge. However, despite its numerous pitfalls, proteomics know-how and methods largely rely on imputation, as realistic complete data are necessary to many quality control methods relying on visualization techniques, clustering, or descriptive data analysis (Webb-Robertson *et al.*, 2015). This is why, we propose here a method to diagnose the missing data mechanism behind each missing value. Moreover, we describe original imputation strategies which specifically account for the nature of each missing value.

In Sec. 2, we introduce notations and data assumptions. In Sec. 3, an original estimator of the MCAR proportion is presented; on its basis, we estimate the posterior probability that each missing value is either MNAR or MCAR. In Sec. 4, we build multiple imputation strategies combining MCAR- and MNAR-devoted algorithms, which are evaluated in Sec. 5.

## 2 Notations and assumptions

### 2.1 Notations

Within each peptide-level dataset, different biological conditions are compared together, by means of several replicated samples (classically, between 3 and 10 samples per condition) so as to account for biological and measurement variabilities. This leads to a matrix with few observations (the replicated samples in each conditions) and several thousands of variables (the union of all the peptides identified in all the samples).

Hereafter, the data structure is a matrix with *n* identified peptides as rows, and 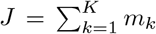 intensity measures for each identified peptide as columns, where *K* is the total number of biological conditions and *m_k_* is the number of samples in each biological condition. In practice, *n* is expected very large compared to *J*, and *K* ≥ 2. However, as this article focuses on imputation, and as it does not make sense to borrow information between different biological conditions to improve the imputation, it simplifies the statistical exposure to consider here that the data matrix contains a single condition (*K* = 1).

For a given sample *j* ∈ [1, *J*], let *F_j_* be the cumulative distribution function (cdf) of the complete intensity values and *π_na_j__* the proportion of missing values among the *n* peptide intensities. Then:

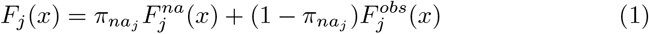

where 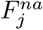 corresponds to the cdf of unknown intensities of missing values 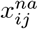, and 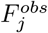 is the one of observed values 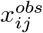.

Moreover, within each sample *j*, MCARs and MNARs coexist in unknown proportions *π_mcar_j__* and 1 − *π_mcar_j__*. As MCARs occur uniformly among the range of intensity levels, their distribution is the same as of complete values. Thus, 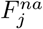 reads:

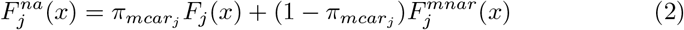

where 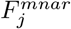 is the MNAR cdf in sample *j*. Eq. (2) can also be written as:

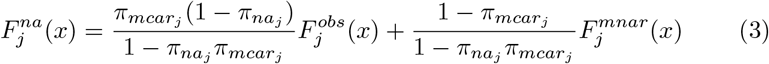

Within Eq. (3), 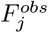 and 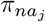 are straightforwardly derived from the data. However, *π_mcar_j__*, 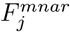 and 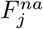 are not, which makes estimating this model impossible without additional assumptions.

### 2.2 Assumptions

Herein, we present some general assumptions regarding the data set which allow the estimation of *π_mcar_j__*. All of them are thoroughly justified in the Supplementary Information.

#### Assumption 1 (Absence of non-quantified peptide).

*Each peptide has at least one observed value among each biological condition.*

#### Assumption 2 (Peptide-wise independence).

*The complete intensity values of peptides are independently distributed in each sample.*

#### Assumption 3 (Intensity distributions).

*(a) The peptide concentrations are log-normally distributed within each sample; (b) the MNARs result of left-censorship which does not impact the most intensely detected peptides.*

Based on these first three assumptions, it is possible to intuitively sketch our strategy to classify each missing value as either MCAR or MNAR:

1. Thanks to Ass. 3b, a subset of peptides with sufficiently high intensity values will only be impacted by MCAR values. This greatly simplifies the estimation task, as it implies that the right hand side of the cdf of missing values (namely 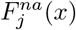 for some large *x*) can be approximated by the cdf of the values imputed by a MCAR-devoted algorithm. Thus, we first impute each missing value 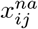 regardless of its nature with an MCAR-devoted imputation algorithm, leading to values which we note thereafter 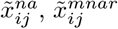 and 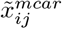 in function of their true nature. We note 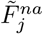 the empirical cdf of the 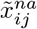.
2. Second, although 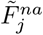 is a rather crude estimate for 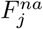 (as there is no reason to expect that the MCAR-devoted imputation will provide unbiased values, especially on lower intensity peptides), it is sufficient to reliably estimate the following quantity:

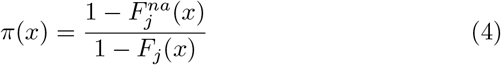

which appears to provide a good estimate of *π_mcar_j__* for some large enough *x*.
3. Third, once an estimate for *π_mcar_j__* is available, it becomes possible to adjust the parameters of *F_j_*, which is a Gaussian (see Ass. 3a). Then, the estimation of 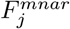 and 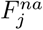 allows to end up with a fully specified model in Eq. (3).
4. Finally, using this model, it is possible to compute the probability that any missing value is MNAR or MCAR, given the intensities of other observed values for the same peptides (which exist, see Ass. 1).

This outline suffers for a single drawback: it appears the estimate for *π*(*x*) has a diverging asymptotic variance at step (1). To cope with this, we make an additional and temporary assumption (see Ass. 4 below) which is a parametric model on the distribution of MNARs for large *x*. This assumption is temporary in the sense that it is only used to stabilize the estimation of *π_mcar_j__* and forgotten right after, so that 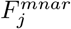 is still a parameter-free distribution at step (3). Before detailing this last assumption, let us formalize some consequences of Ass. 3b:

#### Corollary 1 (Of Ass. 3b).

*Let be*

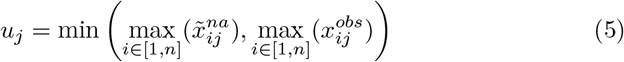

*where* 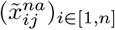 *are the imputed missing values after the use of a MCAR-devoted algorithm. Then*, ∃*M_j_ < u_j_ such that* ∀_*x*_ ≥ *M_j_*:

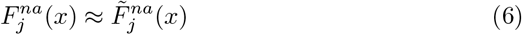

*where* 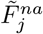 *is the empirical cdf of all the imputed missing values after the use of a MCAR-devoted algorithm.*

#### Corollary 2 (Of Ass. 3b).

*If* 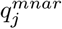 *denotes the theoretical quantile function of MNARs in sample j, then* 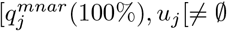.

The proofs of these two corollaries are given in supplemental materials. Finally, our last temporary assumption reads:

#### Assumption 4 (Approximated Weibull cdf of high MNAR values).

∃*M_j_< u_j_ s. t.*

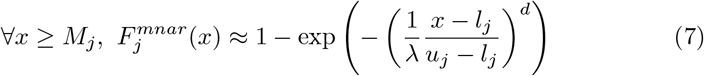

*where d* > 0 *is a shape parameter*, λ > 0 *is a scale parameter*, 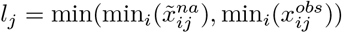 *is an approximation of the minimum of the complete intensity values in sample j, and* 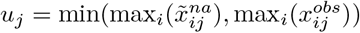 *has been defined in Cor. 3.*

## 3 Estimating *π_mcar_j__* and the nature of each missing value

### 3.1 A first approach to estimate *π_mcar_j__*

Let us consider the following quantity, briefly sketched in Eq. (4):

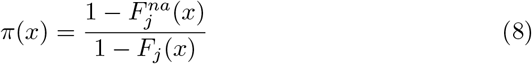

From Cor. 3, 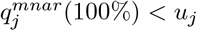, so that 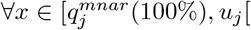

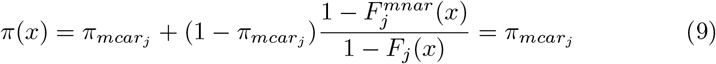

Thus, an estimator of the MCAR proportion derives from an estimate of *π*(*x*) when 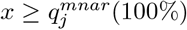. To estimate *π*(*x*), we rely on Prop. 1.

#### Proposition 1.

*Let R and S two independent random variables following, respectively, the binomial distributions* 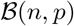 *and* 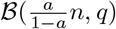 *where* (*a,p,q*) ∈]0, 1[^3^. *We note, respectively, r and s the realizations of R and S. Then, the maximum likelihood estimator (MLE) of θ = q*/(*a × q*+(1 − *a*) × *p*) *is given by 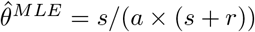 and its asymptotic distribution is*

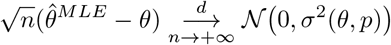

*where the asymptotic variance function is*

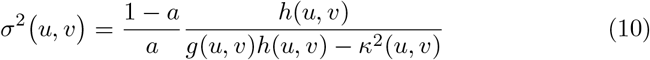

*with*

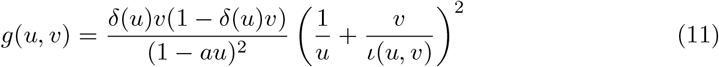

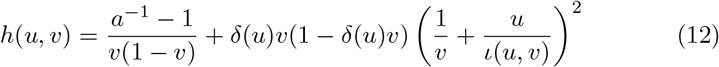

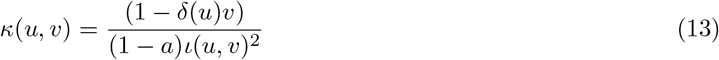

*where* 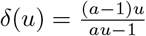 *and* 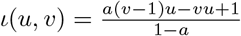.

*Proof.* See Supplementary Information.

Under Ass. 2, 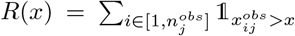 and 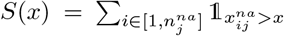 are two independent binomial variables of respective distributions 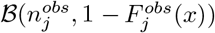 and 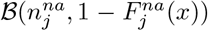 where 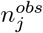 is the number of observed values in the sample *j* and 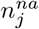 is the number of missing values in the sample *j*. Thus, Prop. 1 provides the MLE of *π*(*x*) (see Supplementary Information for details):

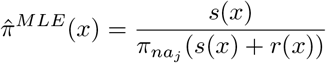

(where *s*(*x*) and *r*(*x*) derive from the empirical cdf). Next proposition shows that, under the Ass. 2 and Cor. 3, an approximation of 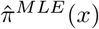 using values imputed by an MCAR-devoted algorithm provides an unbiased estimator of *π*(*x*).

#### Proposition 2.

*Let*

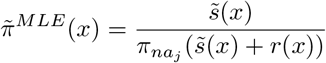

*where* 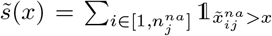 *and* 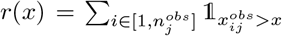. *Under Ass. 2 and Cor. 3, the proportion of missing values π_na_j__ is fixed. Then, for* 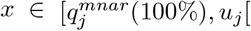,

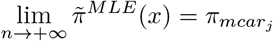

*Proof.* See Supplementary Information.

Although the asymptotic bias of 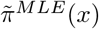 is null when 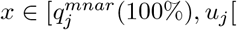 from Prop. 2, the next proposition shows that the variance of 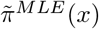 will be high when *x* is close to *u_j_*:

#### Proposition 3.

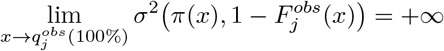

*Proof.* See Supplementary Information.

Hence, if 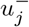 is a maximal value in 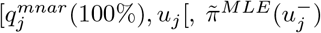 does not seem a wise choice to estimate *π_mcar_j__* in practice. This is why we hereafter rely on a heteroscedastic nonlinear regression to account for this variance in the estimation of the MCAR proportion.

### 3.2 A corrected estimator for *π_mcar_j__* assuming Weibull law

For *x ≥ M_j_* and according to Ass. 4, 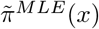 should follow the non-linear regression model defined by:

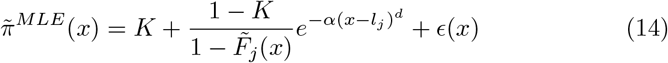

where: *α* corresponds to 1/(λ(*u_j_ − l_j_*))^*d*^ in Ass. 4; 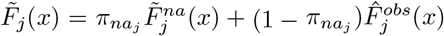 is the empirical cdf of the completed intensity values in the sample *j*; 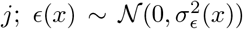 with 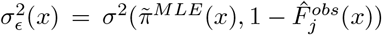; see Eq. (30). Hence, estimators for *K, α* and *d* in Eq. (14) are given by:

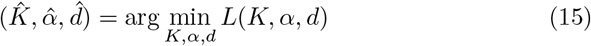

where

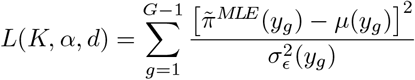

with 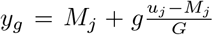 and *G* is a fixed number of sub-intervals of [*M_j_, u_j_*] with equal widths; 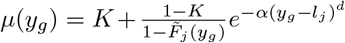. In practice, an appropriate choice is

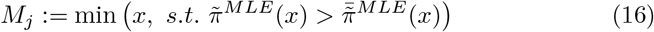

where 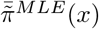 is the average of the 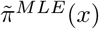 estimated on the interval [*l_j_, u_j_*] (Fig.1A). The variance weighting in the cost function *L*(*K, α, d*) mitigates the impact of the high intensity values in the estimation procedure. A quasi-Newton algorithm with box constraints (Byrd *et al.*, 1995) can be used to minimize Eq. (15) under the following constraints: *K* ∈ [0, 1], *α* ≥ 0 and *d* ≥ 0. Then, according to Eq. (14), *π_mcar_j__* is estimated by 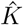.

**Figure 1:**
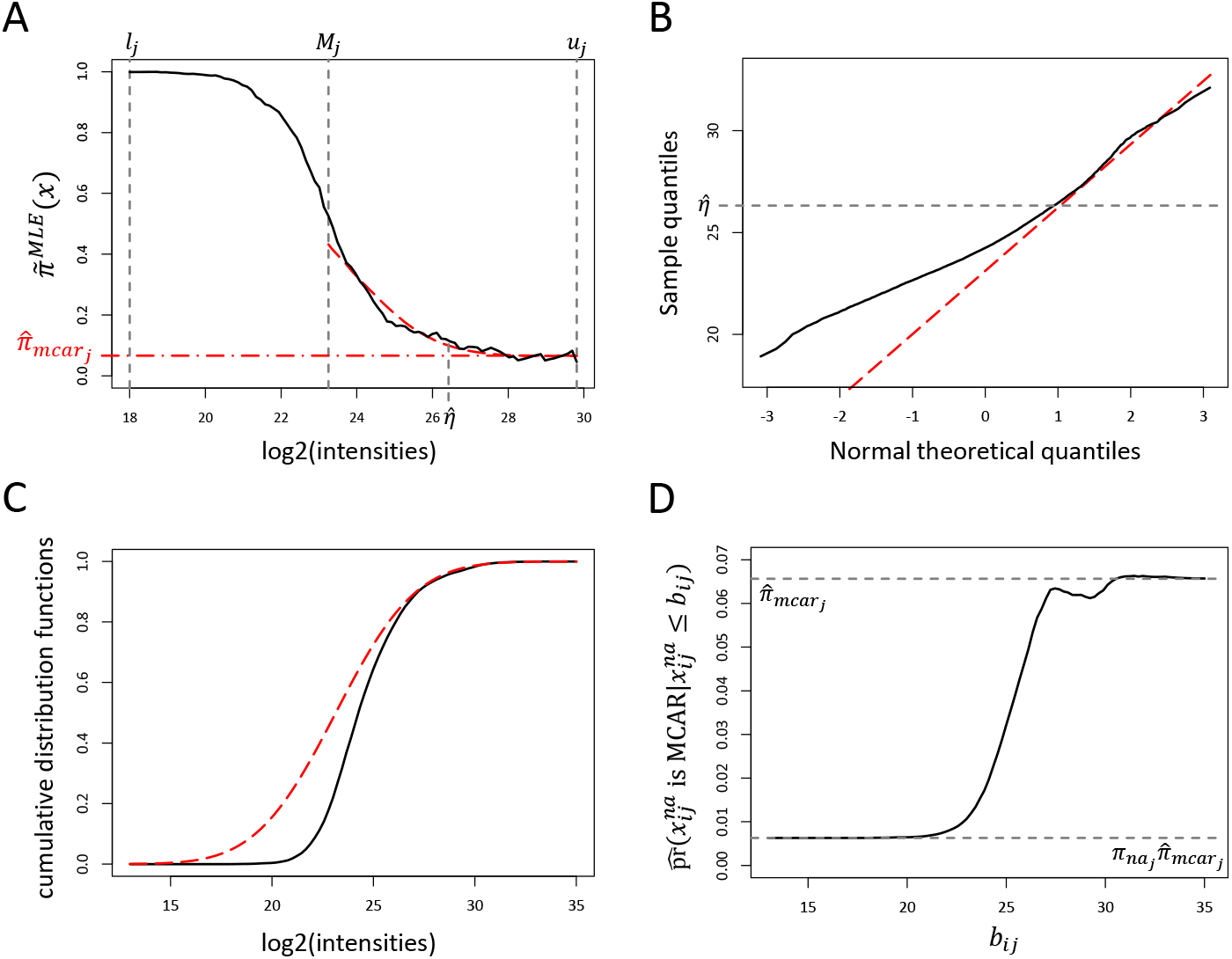
Figure summarizing the different steps of the estimation of *π_mcar_*, and of the probabilities that each missing value is MCAR in a sample of a real dataset (Exp1_R25_pep from the R packages DAPARdata (Wieczorek, 2016; Wieczorek *et al*., 2017)). **A**: the estimated 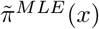 are represented by the black curve, the estimated 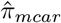 is represented by the red dot-dashed line, while the red dashed line represents the estimated trend (Eq. (14)). **B**: the QQ-plot used to estimate the distribution of complete values. The red dashed line represents the trend line estimated using (Eq. 17). **C**: the cdf of observed values is displayed by the black curve, while the estimated cdf of complete values is the red dashed line. **D**: The estimated probability that a missing value is MCAR (Eq. 21) in function of *b_ij_* in the sample.

### 3.3 Estimation of the distribution of complete values

Once *π_mcar_j__* is estimated, it is possible to estimate the distribution of complete values under Ass. 3. Under this assumption, a straight line must be observed on the Q-Q plot between the quantiles of the observed values and those of a normal distribution, when the quantiles of the observed values become superior to 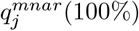 (Fig.1B). To estimate this line, the following regression model is used in each sample *j*:

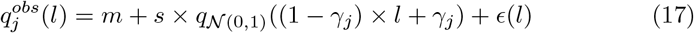

where *ϵ*(*l*) is a Gaussian white noise, 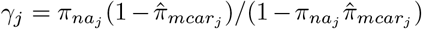, *m* and *s* are the mean and standard deviation of the normally distributed complete values, and 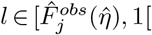 where 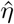 represents a value such that 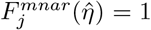. This regression model is motivated by the fact that *F_j_* can be expressed in function of 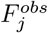 and 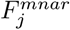 from (1) and (2):

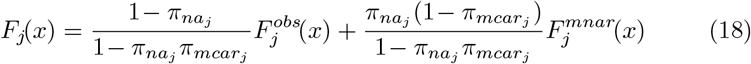

In practice, 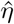 is estimated by searching a minimal value for which the trend of the non-linear model estimated in section 3.2 is no more significantly different from 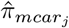:

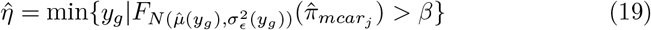

where 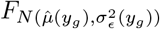 is the cdf of a Gaussian distribution with a trend and a variance equal to the ones of the non-linear model estimated in section 3.2, and *β* is a confidence level, for instance 5%.

### 3.4 Predicting the nature of missing values

Thanks to the estimation of the distribution of complete values, the probability that a given missing value is MCAR conditionally to the fact it is inferior to *b_ij_* can be estimated through the Bayes theorem:

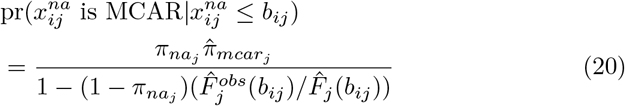

In practice, the intensities of a peptide *i* within a given condition *k* come from replications of a same experiment, so that it makes sense to use a same value as upper bound for all the samples *j* of each condition *k*. In this way, Eq. (20) becomes

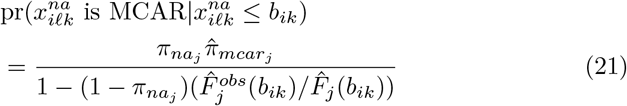

for the *l^th^* sample of the *k^th^* condition. That is why we propose to fix *b_ik_* to the maximum observed intensity value for the peptide *i* in the condition *k*, i.e. 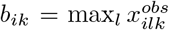. Consequently, the hypothesis that a missing value is MCAR can next be rejected when the probability estimated in Eq.(21) is inferior to a chosen threshold, so that it becomes possible to categorized each missing value as MCAR or MNAR.

## 4 Imputation methods

In literature, many algorithms are available to impute missing values when they are assumed either MCAR or MNAR. For MCAR values, REM (Schneider, 2001) or LS (Bø *et al.*, 2004) algorithms work well on proteomics data according to Webb-Robertson *et al.* (2015). For MNAR values, several methods have been proposed to impute them with small values, since they are mainly assumed to be inferior to the MS detection limit (Deeb *et al.*, 2012). In this context, Lazar *et al.* (2016) showed that the choice of the most suitable method for the MCAR values, or that of the most adapted for the MNAR values, is less important than the choice of applying a method adapted to the true nature of the missing values. We propose here two imputation strategies that allow to adjust the imputation to the nature of the missing values from the probabilities estimated by equation (21).

### 4.1 Naive hybrid imputation strategy

A naive algorithm consists to fix a threshold on the probabilities estimated by Eq. 21 and to impute the missing values according to their estimated nature:

**Algorithm 1.**
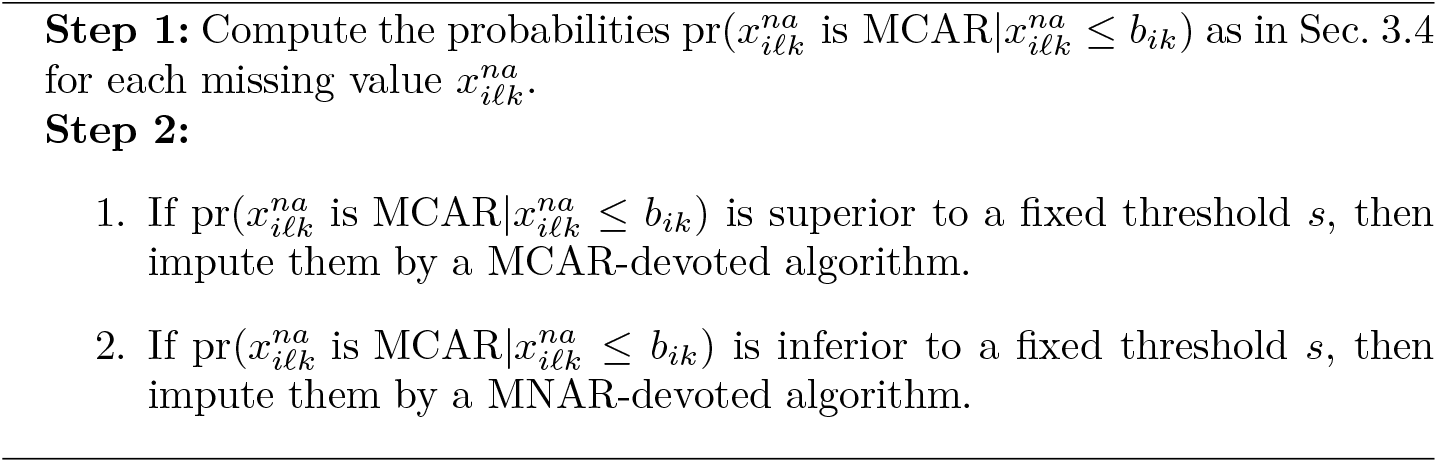
Naive Hybrid Imputation Strategy

This naive approach has two major drawbacks. First, the sample-wise correlations resulting from a possible complex experimental design are not equally accounted for in the MCAR-devoted and MNAR-devoted algorithms: while they are generally considered in MCAR-devoted algorithms, they are not take into account in MNAR-devoted algorithms. Second, the sensitivity to the categorization (which is bound to possible mistakes in addition to the boundary effects) may hinder the quality of the imputation. That is why, we propose an other strategy in the next section.

### 4.2 Multiple imputation strategy

Regarding the drawback concerning the sample-wise correlations of the naive strategy, the only weakness of a good MCAR-devoted imputation strategy is that the distribution used to perform the imputation does not reflect the distribution of MNARs for the simple reason that there are no such observable values to build the distribution on. As a result, it makes sense to use any MNAR-devoted algorithm only to artificially create low intensity values in a first step. Thanks to them, any MCAR-devoted imputation algorithm used afterwards will draw the imputed values by accounting for the multiple natures of missing values. This has a great advantage: it does not matter if the MNAR-devoted algorithm does not account for the experimental design or correlations between samples, as the imputed MNARs are only a preliminary step used to build the final imputation model. Regarding the second drawback concerning the sensitivity of the categorization, it is possible to get rid of the stiff categories by simply assigning each missing value to the MCAR category according to a Bernouilli trial with probability of success pr(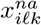 is 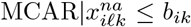) and to perform the imputation accordingly. Then, the process is repeated *N* times. Put together, these ideas naturally lead to the following algorithm:

**Algorithm 2.**
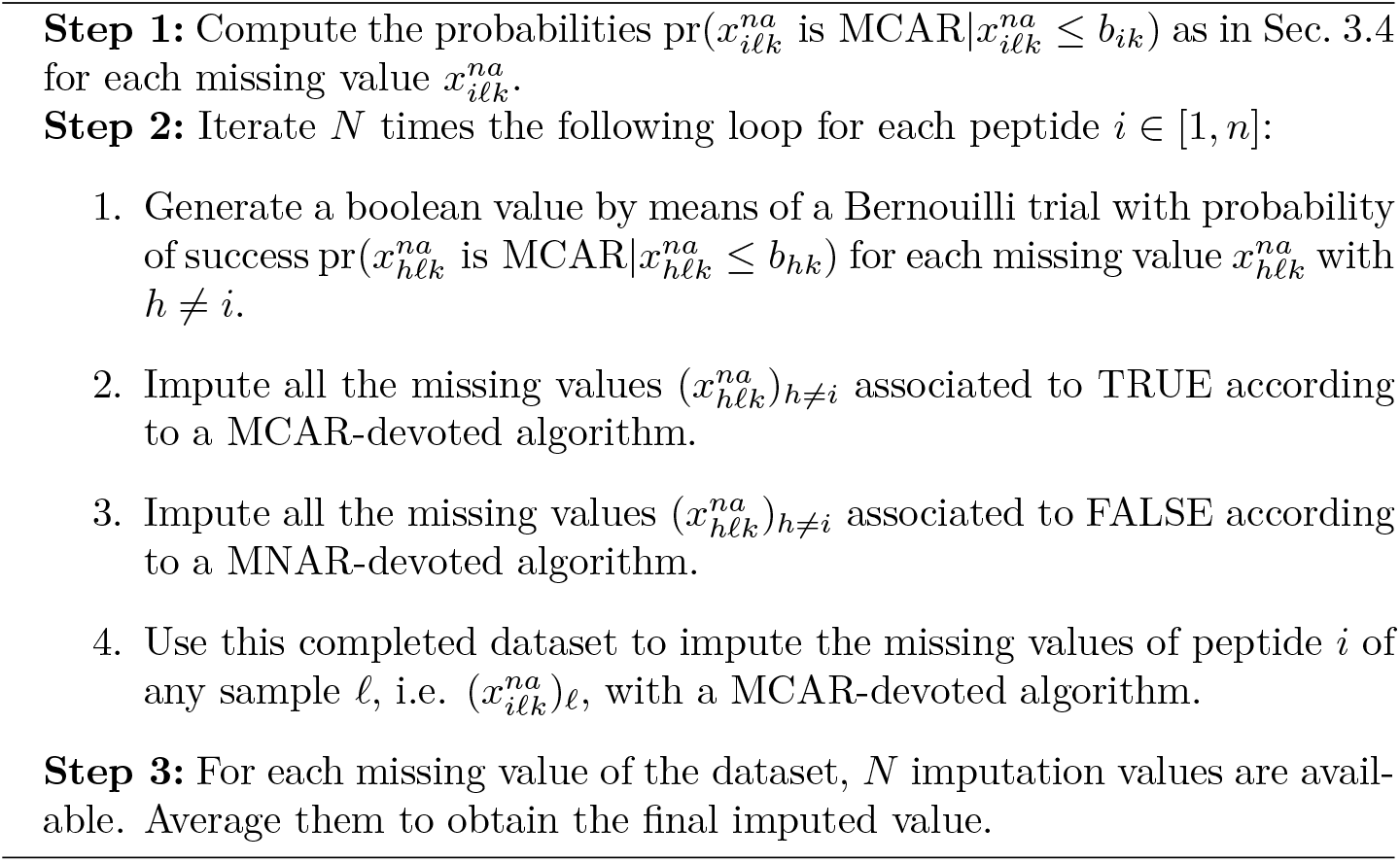
Multiple Imputation Strategy

Possibly, confidence intervals can be obtained by using the Rubin’s rules (Royston *et al.* (2004)). Simulations (see Supplementary Information) suggest that *N* = 10 is sufficient to reach nearly optimal performances. Practically, it is possible to reduce the computational load by calling the MCAR-devoted algorithm only *n × N* times. To do so, one uses the same Bernouilli trials (step 2.1.) and the subsequent imputations (steps 2.2 and 2.3) within each loop.

## 5 Simulation studies

Herein, extensive simulations were used to assess our methodology to diagnose MCAR and MNAR values.

### 5.1 Experimental design

To generate artificial but realistic datasets, we consider a case where *n* peptides were previously identified and quantified in *nc* different biological conditions. Within each condition, there are *nb* different biological samples. For each of the biological samples, *ns* technical replicates are considered. As a result, each data matrix is made of *n* rows and *ns × nb × nc* columns.

#### 5.1.1 Generating complete data

The logarithmized intensity values for the resulting *ns × nb × nc* samples are simulated to mimic their respective LC-MS/MS analysis. For each peptide *i* belonging to the sample *j* coming from the biological sample *b* in condition *k*, logarithmized intensity values *x_ijbk_* are generated from Gaussian distributions 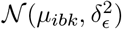 where *μ_ibk_ = μ_ik_ + μ_ib_* with (*μ_ik_*)_*i*∈[1,*n*]_ are independently generated from Gaussian distributions 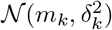 and (*μ_ib_*)_*i*∈[1,*n*]_ are independently and identically distributed following a Gaussian distribution 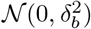.

Concretely, we used the following tuning in our simulations: *n* = 10000, *nc* = 2, *nb* = 3, *ns* = 5, *m_k_* = 25, *δ_k_* = 2, *δ_b_* = 0.5 and *δ_ϵ_* = 0.2 (for it leads to classically observed data; see for example Ramus *et al.* (2016)).

#### 5.1.2 Incorporation of MCAR and MNAR values

Missing values are generated as MCAR values by uniform random drawings without replacement across the list of peptides in each sample. The rest of missing values (i.e. MNAR values) are selected on the basis of random drawings without replacement with the following probability:

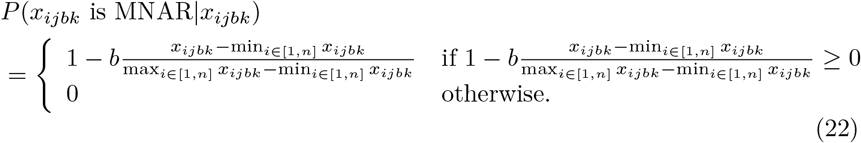

where *b* ≥ 0 allows to adjust the distribution of MNAR values (see Figure 2 for a comparison of several sceniaros with different values for *b*), so that they are distributed more or less similarly to MCAR data. For instance, if *b* = 0, then MNAR values are uniformly distributed among the intensity levels, such as it will be impossible to estimate their proportion and distinguish them from MCAR values. Inversely, the higher *b* is, the easier it is to distinguish them.

**Figure 2:**
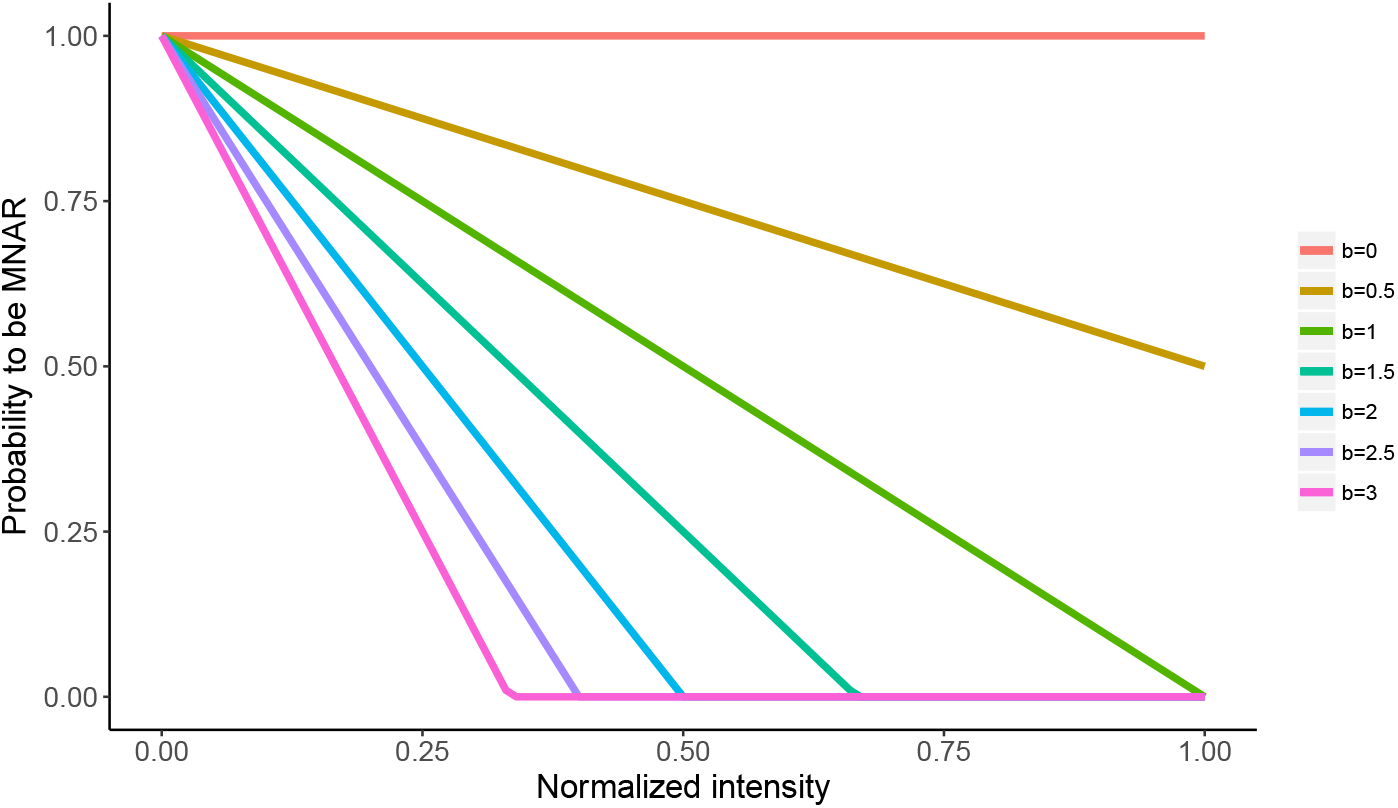
Probability that the missing value is MNAR in function of the normalized intensity level ([*x_ij_ − min*(*x_ij_*)]/[*max*(*x_ij_*) − *min*(*x_ij_*)]) following eq.(22) to incorporate missing values in simulated data.

According to our framework, peptides with which values are all missing in a given condition are removed. This slightly changes the amount of missing values and thus, the proportions of MCAR and MNAR values in each sample: as a consequence, the true values for *π_na_j__* and *π_mcar_j__* need to be re-computed thereafter. All the remaining entries of the data matrix correspond to observed values. This data generation framework is implemented in the R function *sim.data* of the R package *imp4p* Giai Gianetto (2020).

### 5.2 Empirical bias and variance of the estimator of the proportion of MCAR values *π_mcar_*

First, we focus on the quality of the proposed method to estimate the proportion of MCAR values. Concretely, we discuss its empirical bias and variance on data simulated with the aforementioned protocol. Table 1 shows its empirical biases and variances in function of different values of:

- the proportion of missing values *π_na_*,
- the proportion of MCAR *π_mcar_*,
- the *b* parameter, which allowing to adjust the distribution of MNAR values more or less close to the one of MCAR values.

**Table 1:** Empirical biases and variances (in brackets) of 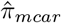 from datasets simulated with different values of *π_mcar_* and *b*.

|  |  |  |  |  |  |  |
| --- | --- | --- | --- | --- | --- | --- |
| $\pi_{na} = 10\%$: | $b$ | | | | | |
|  | 0.5 | 1 | 1.5 | 2 | 2.5 | 3 |
| $\pi_{mcar} = 10\%$ | 8.9% [3.8%] | 20.6% [7.9%] | 10.6% [1.3%] | -1.3% [0.2%] | -3.1% [0.2%] | -4.5% [0.3%] |
| $\pi_{mcar} = 20\%$ | -2.2% [1.6%] | 10.8% [4.4%] | 10.1% [1.3%] | 0.7% [0.6%] | -0.8% [0.5%] | -3.1% [0.6%] |
| $\pi_{mcar} = 30\%$ | -12.8% [1.9%] | 1.1% [1.8%] | 7.1% [1.1%] | 1.6% [0.8%] | 0.6% [0.5%] | -1.5% [0.5%] |
| $\pi_{mcar} = 40\%$ | -23.4% [4.2%] | -6.8% [1.6%] | 1.9% [1.1%] | 0.7% [0.7%] | -0.9% [0.5%] | -2.5% [0.5%] |
| $\pi_{mcar} = 50\%$ | -33.8% [8.6%] | -16.3% [2.7%] | -2.7% [1.6%] | -3.7% [1.1%] | -4.3% [0.9%] | -5.7% [1.0%] |
| $\pi_{mcar} = 60\%$ | -45% [13.8%] | -24.3% [5.4%] | -7.3% [2.6%] | -7.7% [2.2%] | -7.8% [2.1%] | -9.3% [1.9%] |
| $\pi_{na} = 20\%$: | $b$ | | | | | |
|  | 0.5 | 1 | 1.5 | 2 | 2.5 | 3 |
| $\pi_{mcar} = 10\%$ | -0.9% [0.7%] | 10.1% [2.6%] | 8.5% [1.2%] | 3.4% [0.3%] | -1.3% [0.2%] | -4.4% [0.4%] |
| $\pi_{mcar} = 20\%$ | -9.3% [0.8%] | 6.8% [1.2%] | 9.1% [1.3%] | 3.2% [0.6%] | 2.6% [0.3%] | -1.2% [0.3%] |
| $\pi_{mcar} = 30\%$ | -18.5% [2.9%] | -10.4% [0.9%] | 8.4% [1.3%] | 3.9% [0.6%] | 3.3% [0.3%] | -1.4% [0.3%] |
| $\pi_{mcar} = 40\%$ | -28.5% [8.1%] | -18.3% [2.8%] | 4.1% [0.9%] | 3.1% [0.7%] | 1.3% [0.5%] | -2.5% [0.6%] |
| $\pi_{mcar} = 50\%$ | -39.1% [15.7%] | -29.2% [6.7%] | 0.1% [1.5%] | -0.6% [1.1%] | -1.5% [0.9%] | -3.7% [1.1%] |
| $\pi_{mcar} = 60\%$ | -50.1% [26.5%] | -37.9% [11.8%] | -4.7% [2.2%] | -5.2% [1.9%] | -5.8% [1.8%] | -6.8% [1.9%] |
| $\pi_{na} = 30\%$: | $b$ | | | | | |
|  | 0.5 | 1 | 1.5 | 2 | 2.5 | 3 |
| $\pi_{mcar} = 10\%$ | -4.2% [0.4%] | 3.6% [0.9%] | 11.1% [1.8%] | 2.2% [0.4%] | -2.2% [0.3%] | -3.1% [0.3%] |
| $\pi_{mcar} = 20\%$ | -13.8% [1.8%] | -1.1% [0.8%] | 11.2% [1.8%] | 6.8% [0.9%] | 2.2% [0.3%] | -0.2% [0.3%] |
| $\pi_{mcar} = 30\%$ | -21.9% [4.7%] | -17.1% [2.9%] | 10.1% [1.6%] | 7.6% [1.1%] | 2.8% [0.5%] | -0.3% [0.4%] |
| $\pi_{mcar} = 40\%$ | -30.1% [9.8%] | -28.1% [5.4%] | 6.6% [1.5%] | 5.2% [1.2%] | 2.1% [0.9%] | -2.3% [1.1%] |
| $\pi_{mcar} = 50\%$ | -40.3% [17.3%] | -36.6% [11.3%] | 1.6% [1.3%] | 2.1% [1.3%] | -1.8% [1.3%] | -4.8% [1.5%] |
| $\pi_{mcar} = 60\%$ | -51.2% [28.3%] | -44.5% [17.4%] | -2.9% [2.1%] | -2.6% [1.8%] | -3.6% [2.1%] | -7.6% [2.4%] |

For each combination of these 3 parameters, 100 datasets have been simulated.

The results show that our estimator has a weak bias when *b* is greater than 2, that is, when the distribution of MNAR values is shifted far enough to the left. This is in line with the assumptions on which our methodology is based.

However, when *b* is smaller (notably in the cases where *b = 0.5* or *b = 1*), the estimator appears rather imprecise. This is clearly due to the non-respect of Ass 3b. At this point it is important to understand that such imprecise estimation is not really a problem: First, Ass 3b being realistic, datasets are not expected to deviate from it. Second, if a dataset does not respect this assumption notwithstanding, it would not be a problem: in such a dataset, the main difficulty of proteomic imputation, that is the co-existence of two types of missing values with different behaviors has vanished: the missing values that were expected to be MNAR almost behave as MCAR, so that using a single imputation algorithm, such as already proposed in the literature, is sufficient.

Conversely, for simulated datasets with *b* tuned according to what is classically observed (i.e. somewhere between *b = 2.5* and *b = 3*), the measured bias is weaker. More precisely, it seems that our estimator tends to slightly underestimate the proportion of MCAR values when it tends to be very low (*π_mcar_* = 10%), or on the contrary, very high (> 50%). However, these are extreme cases that are not often met on peptide-level data.

### 5.3 Classification of missing values as MCAR or MNAR values

In this section, we only evaluate the capability of discriminating MCAR and MNAR values on the basis of the estimated probabilities of being MCAR (see Eq.21). For each missing value, the prediction is simply made by comparing this probability to a threshold. If the probability is below it, then the corresponding missing value is predicted as MNAR. True and false positive rates are calculated as follows:

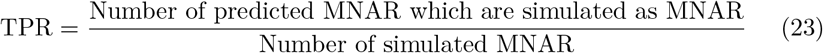

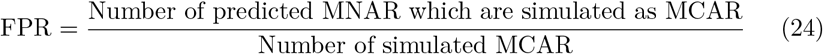

By varying the threshold between 0 and 1, we computed ROC curves (see Fig. 3). Then, the Area Under the Curve (AUC) can be computed (see Table 2). Note that *AUC* > 0.5 indicates our method is more accurate than random classification.

**Table 2:**
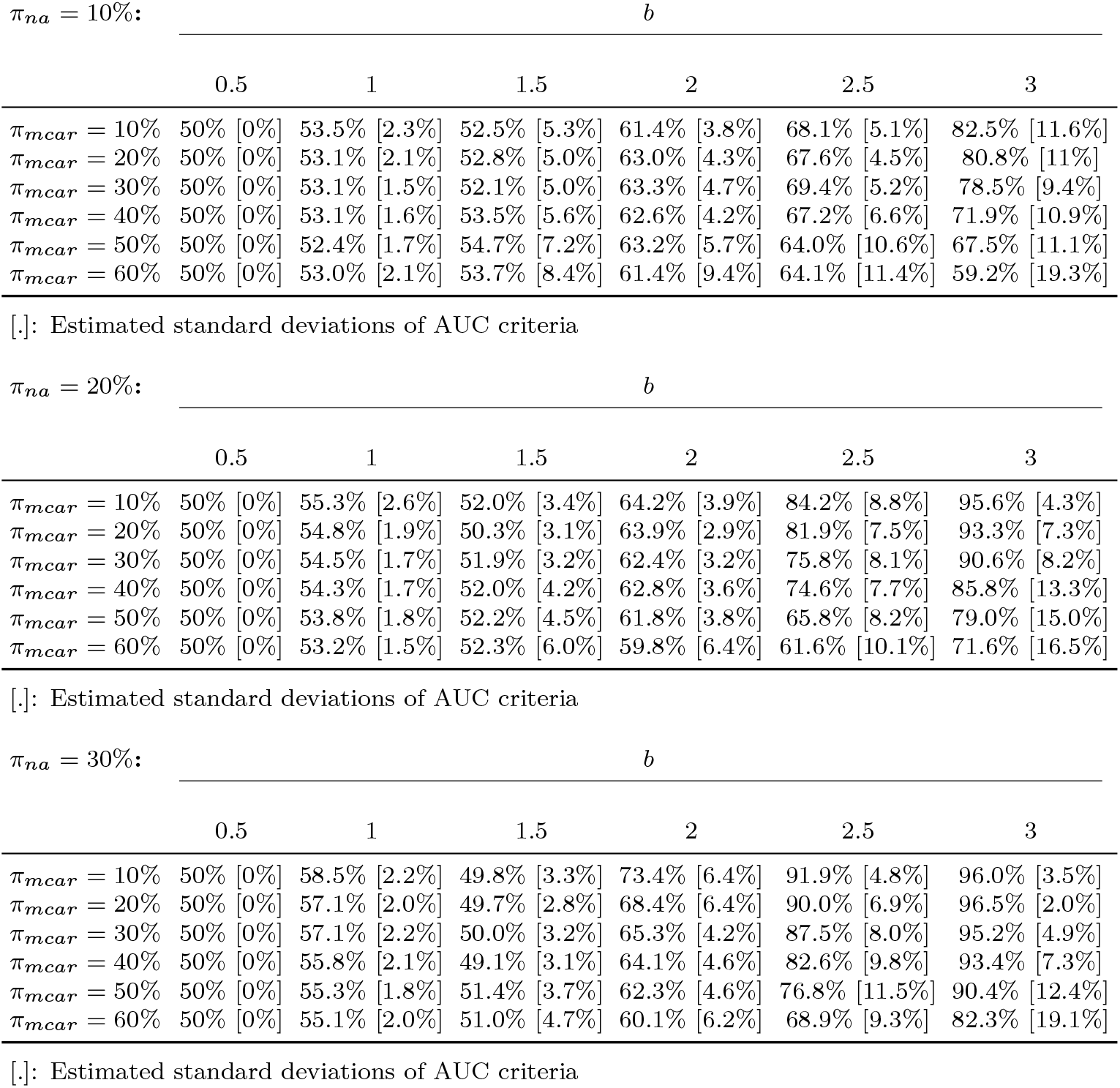
Estimated AUC criteria from datasets simulated with different values of *π_mcar_* and *b*.

**Figure 3:**
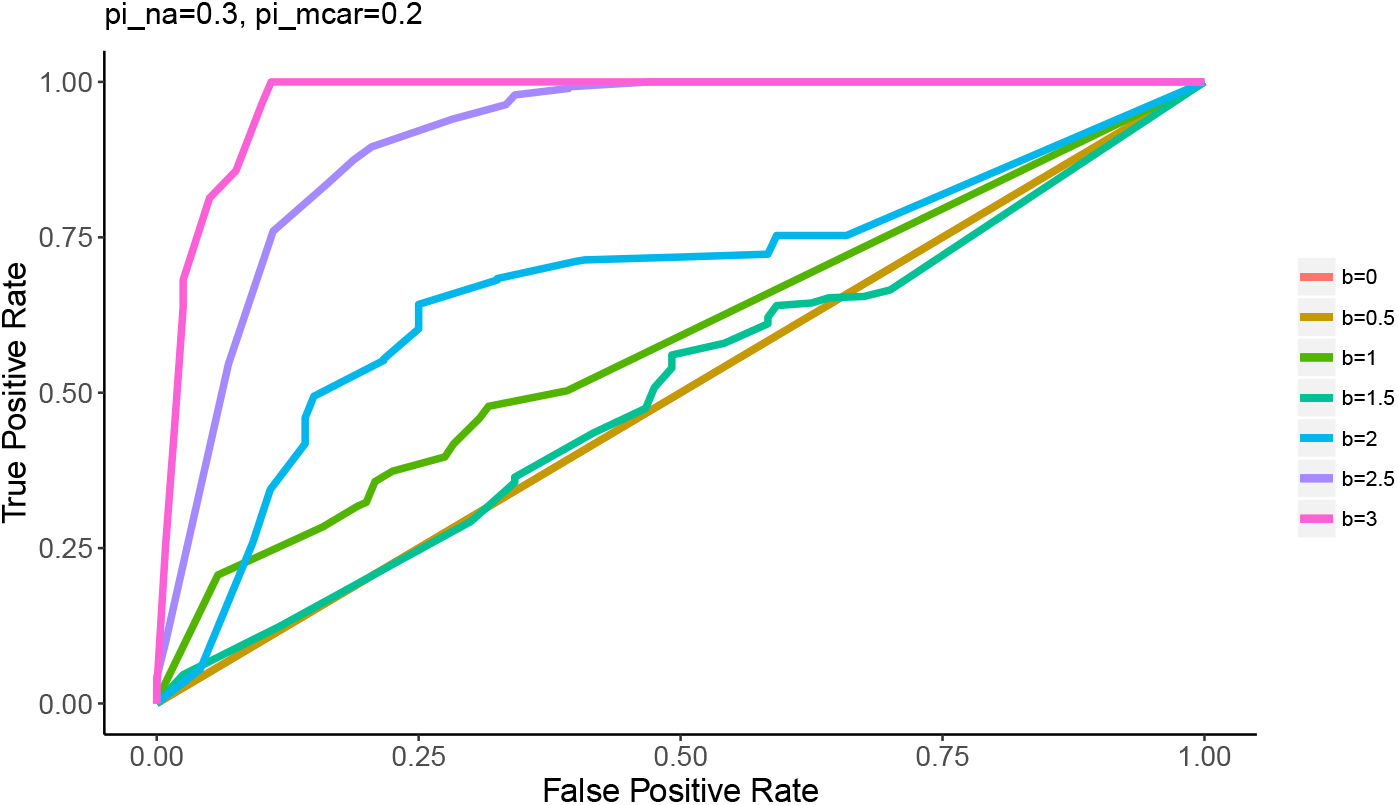
Example of ROC curves obtained with *π_na_* = 30% and *π_mcar_* = 20% for different values of *b*.

From these results, it appears that the AUC criterion is close to 0.5 when the parameter *b* is less than 2, which makes sense, as with such parameter, the missing value generating model is almost the same for MCARs and MNARs, which deviates from our original assumptions. However, the AUC is much better for greater values of *b*. More precisely, it appears the higher *b* is (i.e. the more the distribution of MNAR is shifted to the left) the more reliable the predictions are.

## 6 Comparison of imputation algorithms

In this section, our goal is to compare the proposed imputation methods with other imputation methods. For this, we used the same simulated datasets as in our previous section (see Section 5.1).

### 6.1 Evaluation protocol

#### 6.1.1 Methods and algorithms

In the next, we compare the following algorithms:

- The imputation strategy presented in the article, which calls either SLSA or MLE if the estimated probability of being MCAR is greater than 0.5, and which calls IGCDA otherwise. This algorithm is implemented through the *impute.mix* function of the R package *imp4p* Giai Gianetto (2020). These algorithms are referred to as MIX(SLSA,IGCDA) and MIX(MLE,IGCDA) respectively.
- The multiple imputation strategy presented in the article, which combines either the SLSA or MLE algorithm (as MCAR-devoted algorithm) and the IGCDA (as MNAR-devoted algorithm). The number of iterations for the multiple imputation is discussed next. This algorithm is implemented through the *impute.mi* function of our R package *imp4p* Giai Gianetto (2020).These algorithms are referred to as MI(SLSA,IGCDA) and MI(MLE,IGCDA) respectively.

These methods are evaluated against several MCAR-devoted algorithms:

- The *k*-NN algorithm from Hastie *et al.* (1999), thanks to the *impute.knn* function of the R package *impute* of Hastie *et al.* (2016).
- The BPCA algorithm from Oba *et al.* (2003), thanks to the *pca* function of the R package *pcaMethods* of Stacklies *et al.* (2007).
- The MLE algorithm (described in Section 5.4.1 of Schafer (1997)), thanks to the *imp.norm* function of the R package *norm* of Novo (2013).
- The SLSA algorithm, thanks to the *impute.slsa* function of our R package *imp4p* of Giai Gianetto (2020).

As well as against the following MNAR-devoted algorithms:

- The IGCDA algorithm, thanks to the *impute.igcda* function of our R package *imp4p* of Giai Gianetto (2020).
- The algorithm available in the PERSEUS software Tyanova *et al.* (2016), with default parameters, consisting in imputing the missing values of a sample *j* with small values generated from a Gaussian distribution having a mean equal to 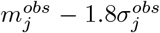 and a standard deviation of 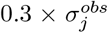, where 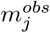 is the average of the observed values in the sample *j* and 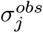 is the standard deviation of these values Deeb *et al.* (2012).

#### 6.1.2 Comparison criteria

To evaluate the discrepancy between the imputed values 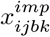 and the ground true values *x_ijbk_*, a Mean Square Error criterion is used:

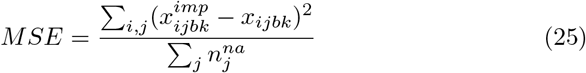

Moreover, to determine if the imputed values has some influence on the variance of a row (i.e. the variance of intensities of a peptide inside a condition), we evaluated the ratio between the variance of the imputed values and the variance of the complete values:

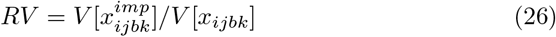

More particularly, as the variance parameter is generally used to perform statistical tests, this criterion allows to evaluate how the imputation methods could impact such tests.

To better visualize these criteria, they are displayed in log scale: As a result, the smaller the log(MSE), the better the method. However, the variance ration RV is expected to have a value as close as 1 as possible, so that log(RV) is not expected to be as small as possible, but on the contrary, to be as close as 0 as possible. Thus, plots depicting log(MSE) and log(RV) should not be interpreted in the same way.

#### 6.1.3 Optimal number of iterations in multiple imputation

When relying on multiple imputation strategies, it is required to tune *N* the number of iterations before the algorithm stops. To estimate it, we simulated 100 datasets with 30*%* of MCAR values among missing values, and we reported the influence of the number of iterations *N* on the quality of the imputation of the *MI*(*SLSA, IGCDA*) algorithm by measuring the *MSE* criterion (Eq. (25)). It appears the *MSE* criterion tends to decrease with *N* (see Figure 4). More precisely, it decreases rapidly between 1 and 3 iterations and seems to reach a plateau after 5 to 7 iterations. In our comparisons with existing algorithms, we set *N* = 10 to garantee optimal performances. Note that these observations are in accordance with the recommendations in Rubin (1987): using between 2 and 10 iterations in multiple imputation procedures are enough to obtain satisfying efficiency.

**Figure 4:**
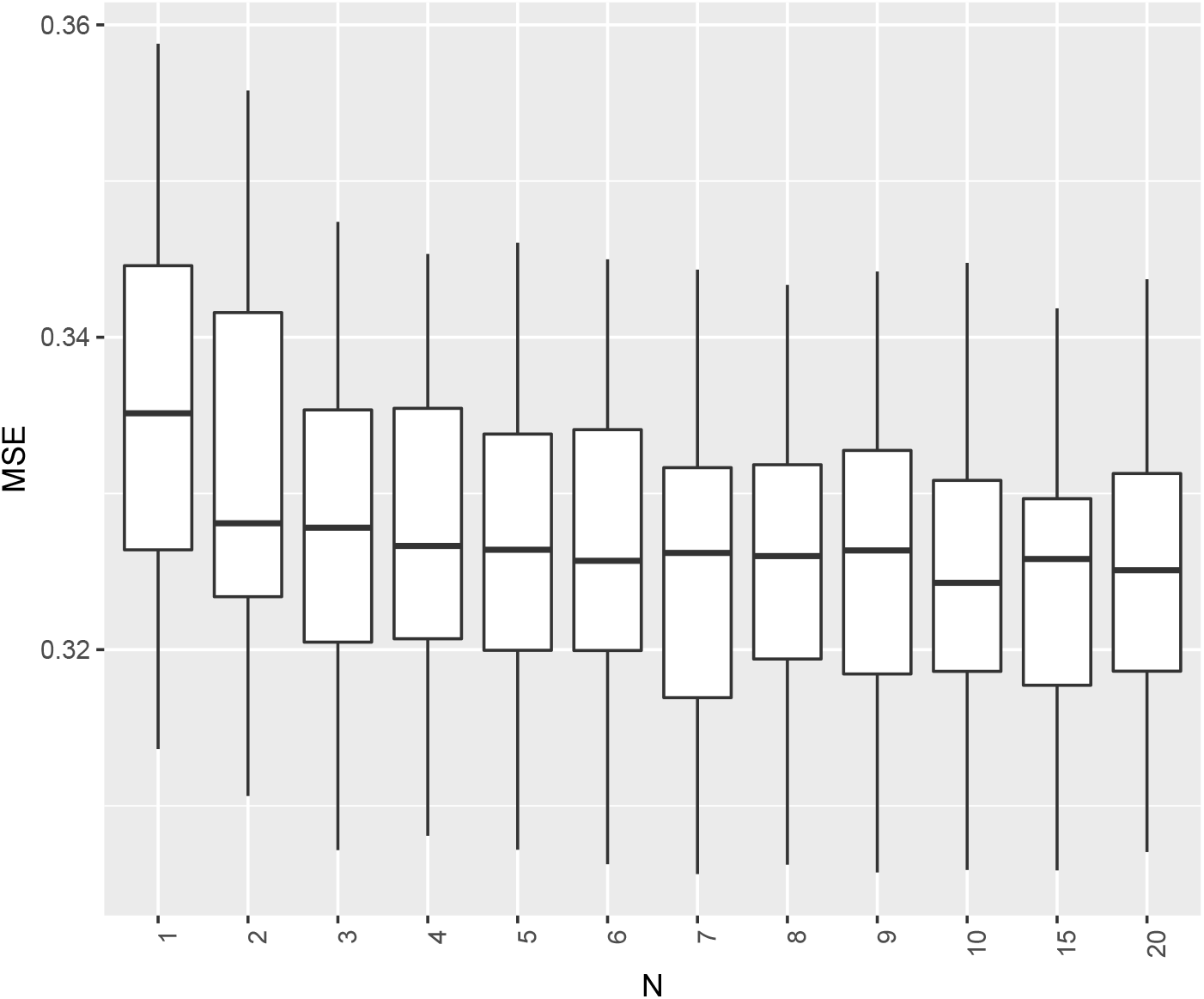
Distribution of the MSE criterion of the algorithm *MI*(*SLSA, IGCDA*) in function of the number of iterations estimated from 100 generated datasets for each iteration.

#### 6.1.4 Comparison scope

To conduct a sensitivity analysis of the different algorithms, we varied three parameters:

- The proportion of missing values *π_na_j__*,
- the proportion of MCAR values *π_mcar_j__*,
- the parameter *b* used to simulate the distribution of MNAR values.

We fixed either *b* = 1.5 or *b* = 3, since *b* = 1.5 represents a case where our methodology is not able to have an AUC criterion different of 50% while *b* = 3 represents a case where our methodology gives an AUC criterion superior to 90% and allows thus a good discrimination of MCAR and MNAR values.

### 6.2 Results

#### 6.2.1 The proposed multiple imputation strategy gives good performances when compared to the other imputation algorithms

From our simulations, the PERSEUS algorithm gives the highest MSE whatever the values of *b*, *π_na_* and *π_mcar_* (see Fig. 5 and Fig. 7). This algorithm imputes missing values with values which are inferior and further from the ground true values than other algorithms. The IGCDA algorithm, which is also a MNAR-devoted algorithm, seems to give a better MSE than PERSEUS. However, it remains quite high when compared to the other algorithms (either MCAR-devoted or those accounting for all types of missing values). These high MSE were expectable, and are due to the principle of imputing all the missing values with small values: as detailed in Lazar *et al.* (2016), this strategy is bound to fail when there are too many MCAR, regardless of the MNAR-devoted algorithm quality. Moreover, these low imputed values will lead to an important increase in the variance of the data and will thus have a strong negative impact on subsequent statistical tests (Fig. 6 and Fig. 8).

**Figure 5:**
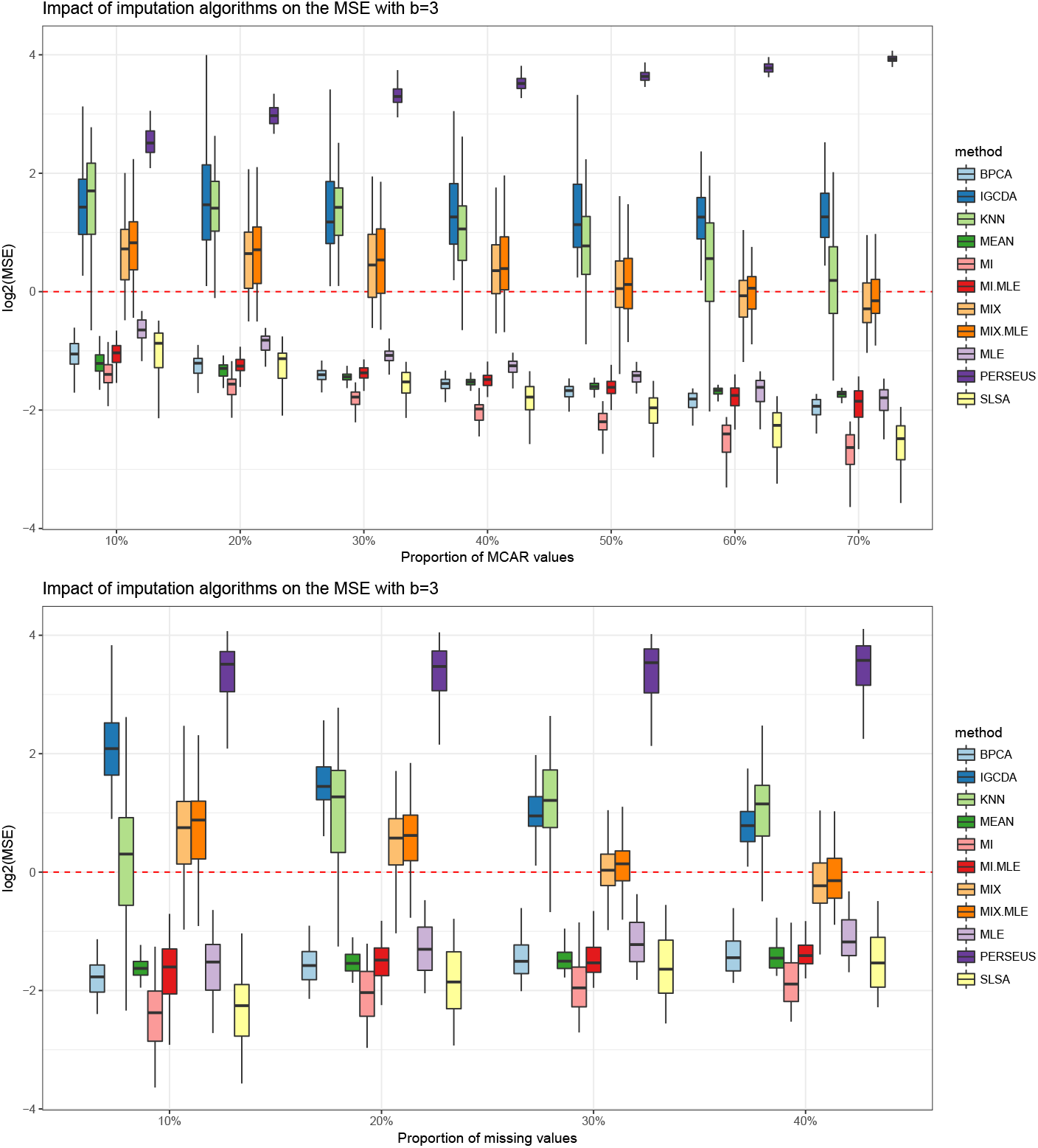
Distributions of the log2(*MSE*) in function of the proportion of MCAR values (top) and in function of the proportion of missing values (bottom) when *b* = 3.

**Figure 6:**
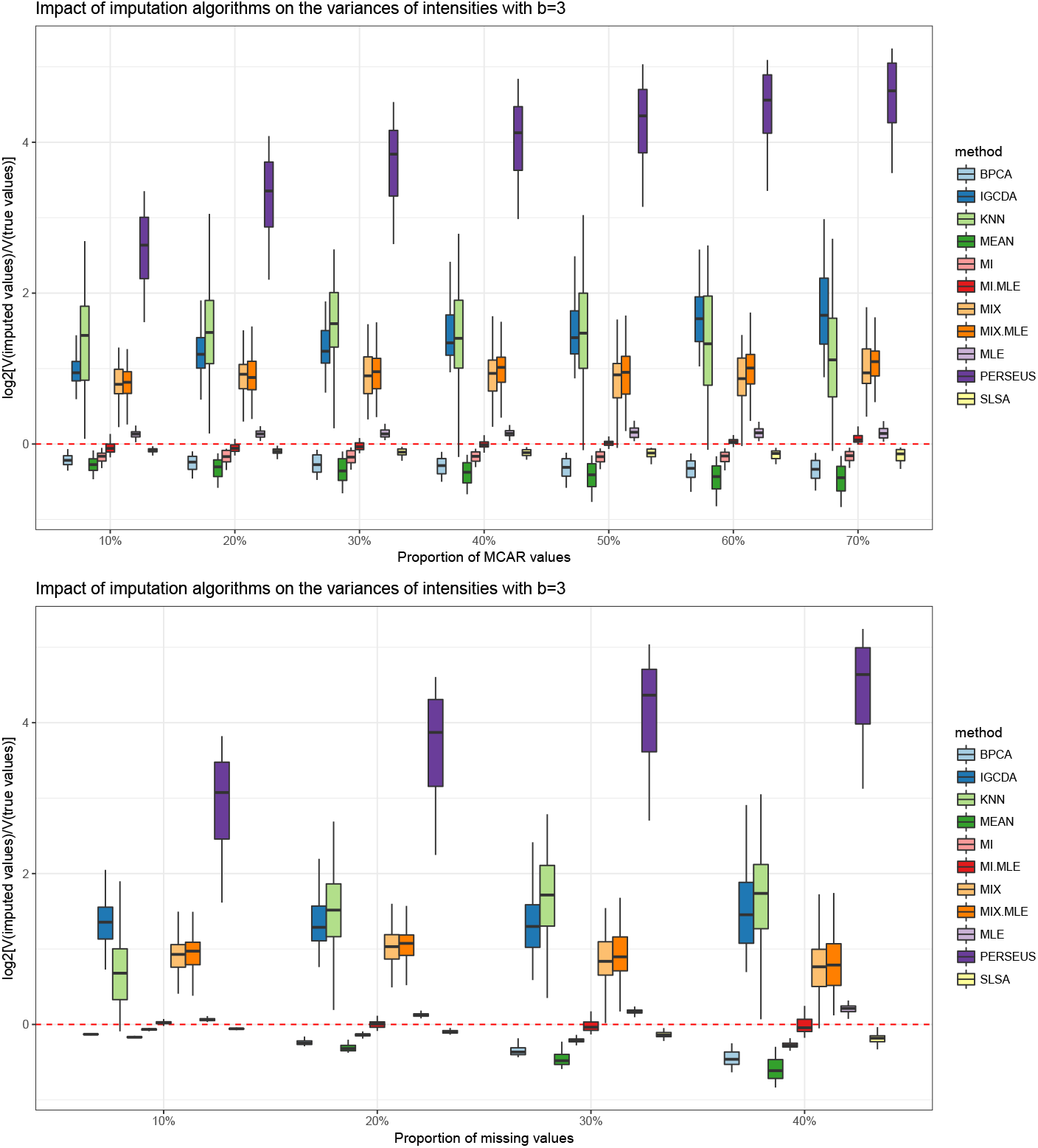
Distributions of the log2(*RV*) in function of the proportion of MCAR values (top) and in function of the proportion of missing values (bottom) when *b* = 3.

**Figure 7:**
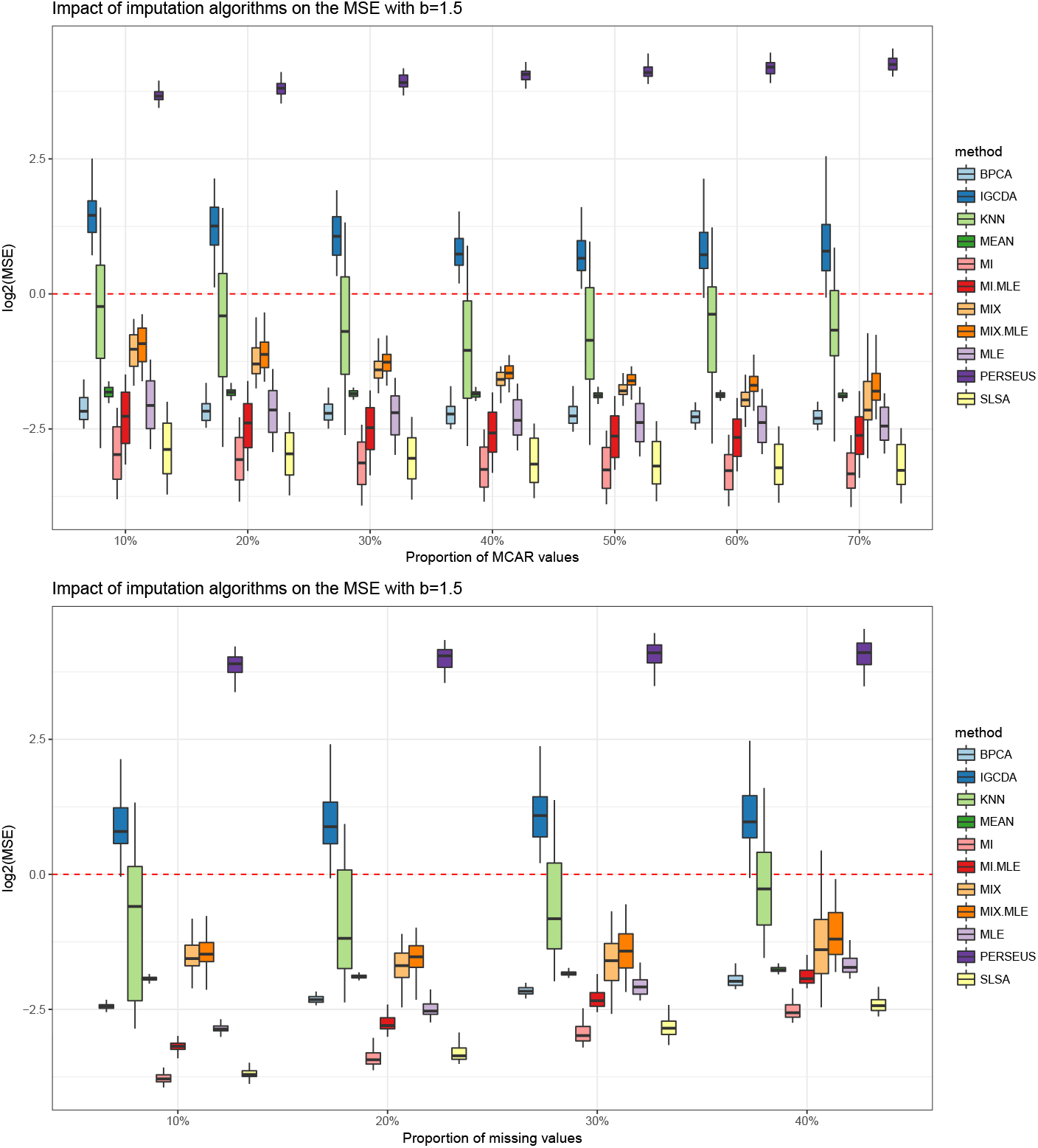
Distributions of the log2(*MSE*) in function of the proportion of MCAR values (top) and in function of the proportion of missing values (bottom) when *b* = 1.5.

**Figure 8:**
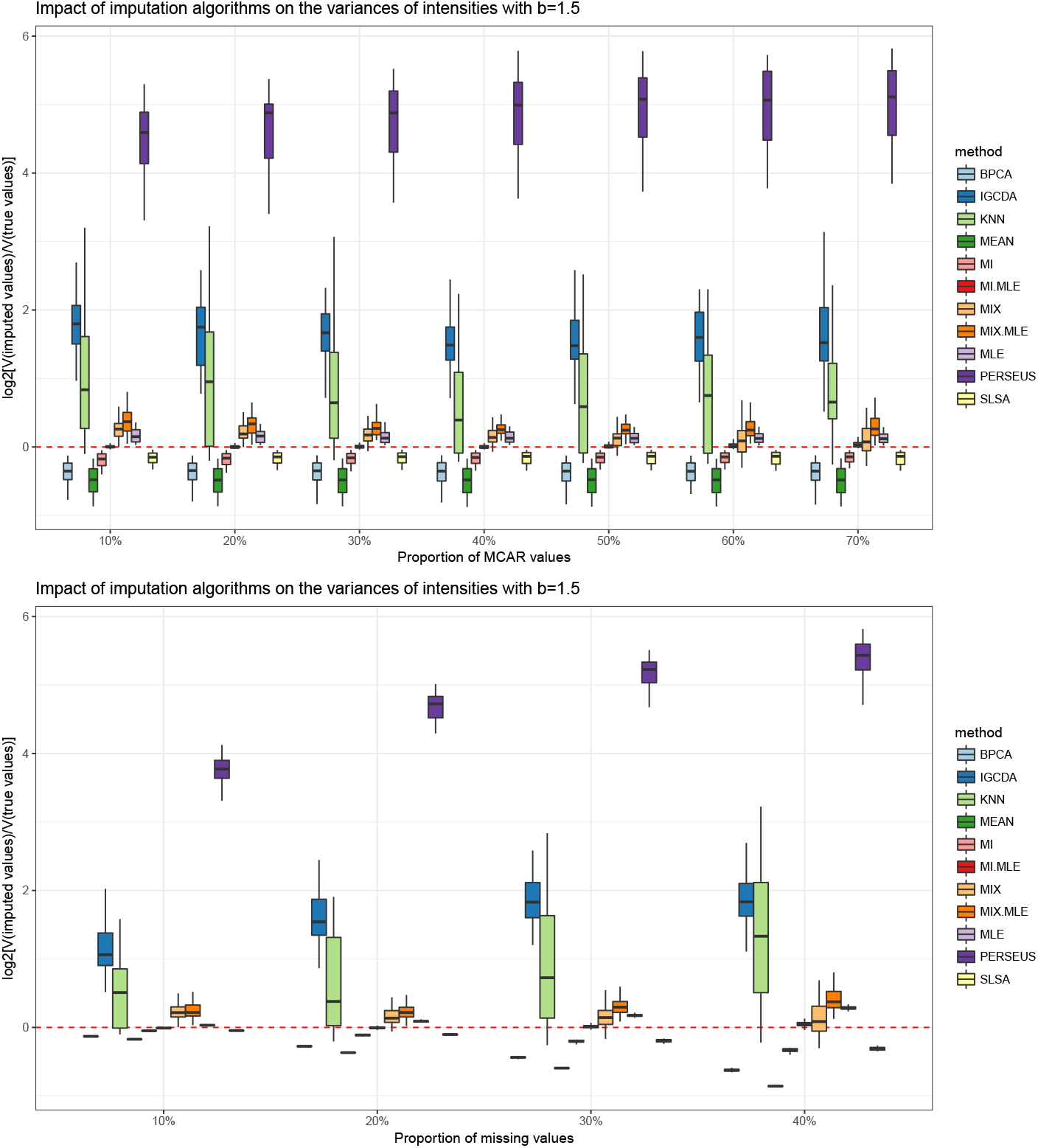
Distributions of the log2(*RV*) in function of the proportion of MCAR values (top) and in function of the proportion of missing values (bottom) when *b* = 1.5.

Another very popular method, namely *k*-NN, gives rather poor results in our simulations. It is the MCAR-devoted algorithm with the highest MSE and it tends to increase the variance of the data (Fig. 5-8). In the case *b* = 3 (the most realistic one), it even gives results equivalent to the IGCDA algorithm (Fig. 5). Moreover, its performances are particularly unstable (it has the widest error bar in the plots of Fig. 5-8).

Two of our proposed strategies, namely MIX(SLSA, IGCDA) and MIX(MLE, IGCDA), also display disappointing results: Although they perform when compared to the aforementioned algorithms (PERSEUS, IGCDA and *k*-NN) they underperform with respect to the MCAR-devoted algorithm they are based on. MIX(SLSA, IGCDA) is less efficient than SLSA, and the same goes for MIX(MLE, IGCDA) and MLE. Globally, it appears that mixing an MCAR-devoted algorithm with an MNAR one to account for MNAR values is not a wise strategy (see Fig. 5-8). This could be extrapolated from the now well-established fact (see Lazar *et al.* (2016)) that in presence of both MNAR and MCAR values, MNAR-devoted algorithm does not perform well with respect to MCAR-devoted ones.

Finally, the remaining algorithms, i.e. BPCA, MEAN, SLSA, MI(SLSA,IGCDA), MLE, MI(MLE, IGCDA) give low MSE criteria when compared to the other algorithms. Among them, the MEAN algorithm has a specific behavior. Unsurprisingly, it is the one providing the lowest RV. In other words, it is the algorithm which most reduces the variance of the data. Although practitioners generally appreciate such type of behavior, as it can somehow compensate for some sources of variability, such type of imputation can artificially increase the number of proteins that appears significantly differentially abundant in the subsequent statistical test, and thus increase the proportion of false positives.

We can also notice a regular pattern in the performances of SLSA and MLE: SLSA always has a better MSE than MLE, while MLE has a better RV than SLSA. This can be observed in any imputation setting: as stand-alone MCAR imputation, within MIX imputation or within MI imputation. BPCA does not perform as well as SLSA or MLE, while being particularly computationally demanding, so that integrating it into MI or MIX strategies does not seems interesting.

Although not very visible, it another trend can be grasped: While MIX imputation strategies did not improve the results, the MI ones are clearly efficient. They always improve the behavior of the associated MCAR method: MI(SLSA, IGCDA) and MI(MLE, IGCDA) perform better than SLSA and MLE respectively, while the difference is more visible with MLE than with SLSA. This will be studied in more details hereafter.

Overall, the algorithm giving the lowest MSE in average whatever the values of *b*, *π_na_* and *π_mcar_* is the multiple strategy MI(SLSA, IGCDA), while the algorithm which seems to change the less the variance of the data is the multiple imputation strategy MI(MLE, IGCDA).

#### 6.2.2 The proposed multiple imputation strategy allow to improve the performance of the MCAR-devoted imputation algorithm on which it is based

As previously sketched, it appears our multiple imputation strategy allows to reach a lower MSE criterion than the MCAR-devoted imputation algorithm on which it is based. This can be easily pictured on Fig. 9. This figure displays, for a given MCAR algorithm **X**, the following ratio:

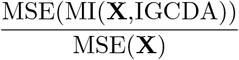

which appears to be in average, constantly below 1. The MSE criterion is even lower as there are MNAR values in the data sets (bottom of Fig. 9).

**Figure 9:**
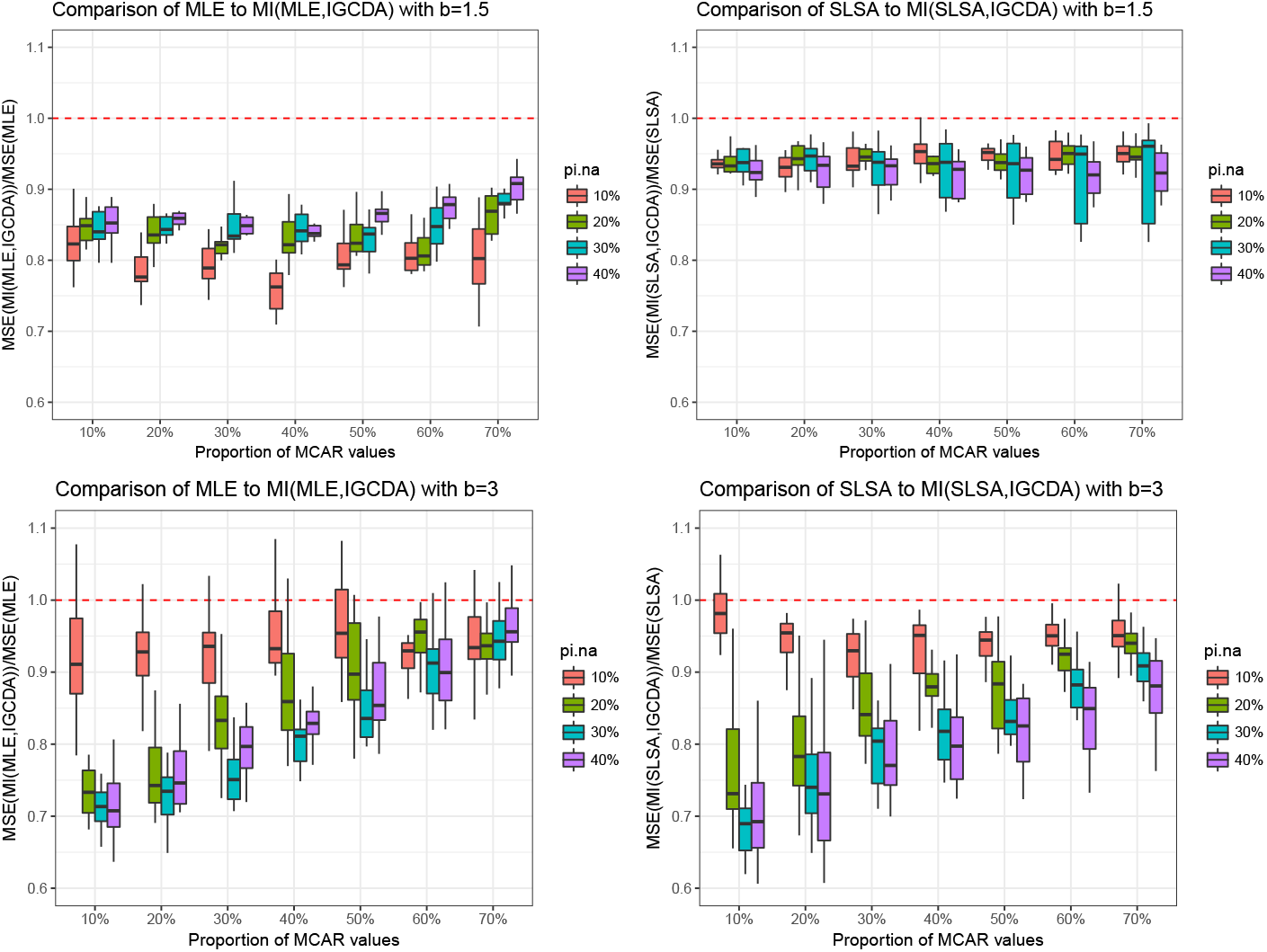
Distributions of the ratios MSE(MI(MLE,IGCDA))/MSE(MLE) (left) and MSE(MI(SLSA,IGCDA))/MSE(SLSA) (right) when *b* = 1.5 (top) and *b* = 3 (bottom).

This feature is more clearly observed when comparing either *SLSA* to *MI*(*SLSA, IGC* or *MLE* to *MI*(*MLE, IGCDA*) when *b* = 3. This makes sense for the following reason: When *b* = 1.5, this feature is not observed since our strategy has difficulties to discriminate between MNAR and MCAR values (AUC close to 0.5%) (top of Fig. 9). However, when *b* = 3, the lower the proportion of MCAR values, the more the MSE criterion is improved using the multiple imputation strategy. Note that this feature is checked only when there is enough missing values, i.e. when *π_na_* > 10% (bottom of Fig. 9). When *π_na_* is too small (case *π_na_* = 10%), the IGCDA algorithm does not allow to shift sufficiently the correlation structure of the dataset; thus, the MCAR-devoted algorithm which is subsequently applied on the imputed values gived results similar to the original MCAR-devoted algorithm.

#### 6.2.3 Conclusion on the comparison of the imputation algorithms

From our simulations, it appears the multiple imputation strategy MI(SLSA, IGCDA) gives the best results in term of accuracy (measured by the MSE criterion) whatever the parameters *b, π_na_* and *π_mcar_* used to generate the data, while the MI(MLE,IGCDA) is the best to preserve the variance of the data. The improvement of the accuracy of the imputations when compared to the SLSA or the MLE algorithm is even stronger that there are MNAR values in the dataset, provided that enough values are missing.

Although our simulations show the relevance of our approach, it has to be noted that the evaluation of a MSE criterion can be subject to cautions. Indeed, the true MNAR values could be far from the ones we simulated, and it is difficult to have an idea of how these values are really distributed in real datasets. However, our simulations are based on a reasonable assumption of a Gaussian distribution of intensities among replicates for a specific peptide. This is reasonable because the use of replications should lead to measured values with a Gaussian-type spread.

As a whole, these simulations clearly stress the gain resulting from our proposed multiple imputation strategy, as well as the underlying diagnosis on the missingness mechanism.

## 7 Conclusion

This article addresses the issue of missing values in peptide-level quantitative proteomics datasets resulting from mass spectrometry experiments. We proposed an approach to estimate the missingness mechanism, as well as a multiple imputation strategy based on this estimation. Our approach assumes a mix of MCAR values and MNAR values, the latter ones being the result of a random left-censorship of the peptide intensities. It allows estimating the proportion of MCAR values and provides an estimate of their distribution among the intensity values of the identified peptides in a sample. We showed it is possible to estimate a probability that each missing value is either MCAR or MNAR. These probabilities can then be used to build multiple imputation strategies combining imputation algorithms dedicated to MNAR values and others dedicated to MCAR values. Our approach is original with respect to the state of the art in the proteomics community where only algorithms based either on a MNAR data assumption or on a MCAR data assumption have been used to date (Lazar *et al.*, 2016). Our evaluations on simulated datasets proved the outperformance of our method with respect to the state of the art, especially when datasets have a large number of missing values. Our methodology could be used in other contexts based on mass spectrometry experiments, such as in metabolomics for example, or, more broadly, on datasets with similar missing value mechanisms. Moreover, the methods and algorithms developed in this article are grouped in a R package named *imp4p*, freely available on the CRAN Giai Gianetto (2020), and wrapped to Prostar software (Wieczorek *et al.*, 2017).

## Supplementary Materials

### 1 Justifications of article assumptions and corollaries

#### Assumption 1 (Absence of non-quantified peptide).

*Each peptide has at least one observed intensity value among the samples of each biological condition.*

*Justification:* Ass. 1 only stipulates that lines from the data matrix that are empty are not considered (obviously, it will not be possible to impute anywhere close to the truth for a peptide in a given biological condition if no observation is available to rely on). This may look like a truism, however, in practice, a peptide may be seen in one biological condition and not in the other, so that from the proteomic practitioner viewpoint, such empty lines exist. We therefore assume they have been previously filtered out and processed separately.

#### Assumption 2 (Peptide-wise independence).

*The complete intensity values of peptides are independently distributed in each sample.*

*Justification:* At first look, Ass. 2 may seem inadequate: it could be naturally expected to observe a strong correlation among intensities of peptides coming from a same protein. However, in practice, there are generally very few proteins with numerous peptides and numerous proteins with few peptides within a sample. Moreover, in practice, several peptides with a similar concentration may lead to measured intensities that differ from several orders of magnitude Silva *et al.* (2005). This oddity comes from the fact that the MS signal of a given peptide is not only influenced by its quantity. It is also strongly dependent on a variety of physicochemical peptide-specific properties, such as its ionization capability. In fact, this is the very reasson why, in absence of isotope labelling (see Introduction), quantitative proteomics is mainly relative: one does not compare the abundance of several peptides within a biological sample, but on the contrary, the abundance of one peptide in samples corresponding to different conditions (leading to assume *K* ≥ 2 in the practical setting described in article, Section 2.1). For all these reasons, the independence assumption of the peptide intensity distribution within each sample is harmless.

#### Assumption 3 (Intensity distributions).

*(a) The peptide concentrations are log-normally distributed within each sample, and (b) the MNAR values result of a left-censorship mechanism which does not impact the most intensely detected peptides.*

*Justification:* These are well-established facts from the literature. For short, Ass. 3a is the reason why quantitative analysis is classically conducted on log-transformed data Eidhammer *et al.* (2012). As for Ass. 3b, some elements were sketched in the Introduction, but a thorougher description of the underlying physico-chemical phenomena can be found in Lazar *et al.* (2016).

#### Corollary 3 (Of Ass. 3b).

*Let be*

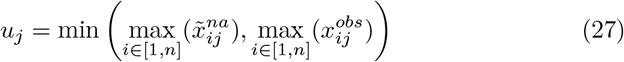

*where* 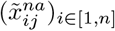 *are the imputed missing values after the use of a MCAR-devoted algorithm. Then*, ∃*M_j_ < u_j_ such that* ∀_*x*_ ≥ *M_j_*:

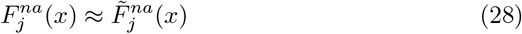

*where* 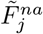 *is the empirical cdf of all the imputed missing values after the use of a MCAR-devoted algorithm.*

*Proof:* 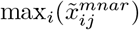 is clearly an acceptable candidate for *M_j_*. Indeed, beyond this value, all the missing values are well imputed by a MCAR-devoted imputation algorithm. Moreover, 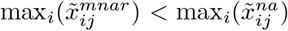 and 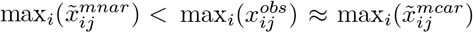 as there exists MCAR values that are beyond the range of left-censored MNAR values. This justifies 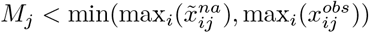.

#### Corollary 4 (Of Ass. 3b).

*If* 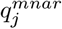 *denotes the theoretical quantile function of MNAR values in sample j, then the interval* 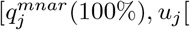 *is non-empty*.

*Proof:* It directly derives from the fact that 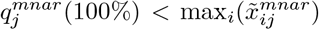 since the overall MCAR-based imputation has overestimated the left-censored values.

#### Assumption 4 (Approximated Weibull cdf of MNAR values).

*It is assumed that* ∃*M_j_ < u_j_ such that* ∀_*x*_ ≥ *M_j_*:

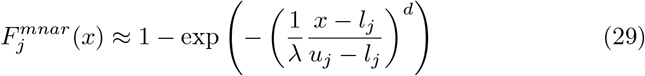

*where d* > 0 *is a shape parameter*, 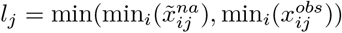 *is an approximation of the minimum of the complete intensity values in sample j, and* 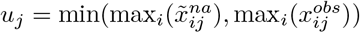 *has been defined in Cor. 3*.

*Justification:* It has to be noted that overall, the distribution of MNAR values may be far from the Weibull distribution. However, this assumption is harmless for at least three reasons: First, one only assume that these two distributions are close when *x* becomes sufficiently great. Second, Weibull distribution is a rather flexible model for left-skewed distributions. Third, one only temporary relies on this parametric model to stabilize *π_mcar_j__*.

## 2 Proofs of the article propositions

### 2.1 Proof of Proposition 1

#### Proposition 1.

*Let R and S two independent random variables following, respectively, the binomial distributions* 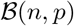 *and* 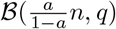 *where* (*a,p,q*) ∈]0, 1[^3^. *We note, respectively, r and s the realizations of R and S. Then, the maximum likelihood estimator (MLE) of θ = q*/(*a × q* + (1 − *a*) × *p*) *is given by* 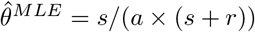 *and its asymptotic distribution is*

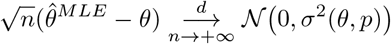

*where the asymptotic variance function is*

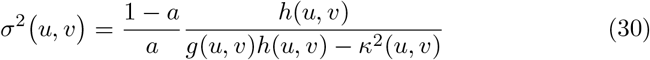

*with*

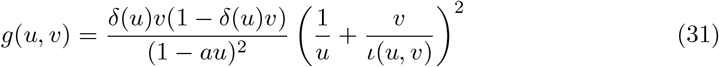

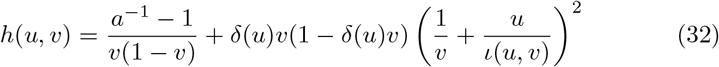

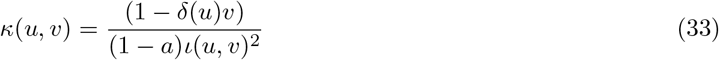

*where* 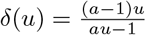 *and* 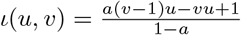.

*Proof:*

**MLE estimates:** With observations *r* and *s* of random variables *R* and *S*, the log-likelihood function reads:

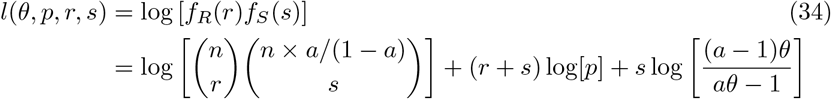

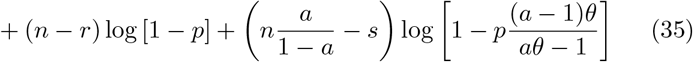

From this log-likelihood function, we have the following MLE of *p* and *θ*:

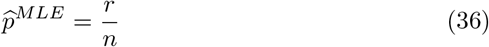

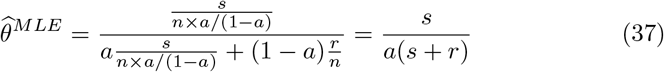

**Fisher Information matrix:** Using the *δ*(.), *g*(., .), *h*(., .) and *κ*(., .) notations introduced in the the statement of Prop. 1, the Fisher information matrix of *θ* and *p* reads:

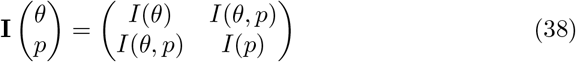

where:

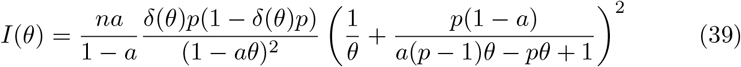

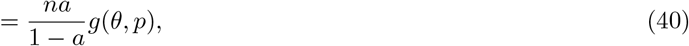

knowing that 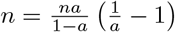:

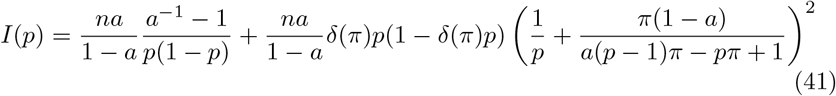

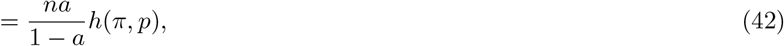

and:

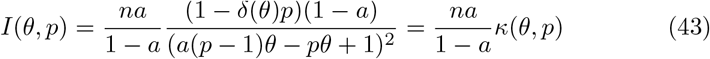

**Asymptotic variance of 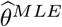:** Therefore, the determinant of the Fisher information matrix reads:

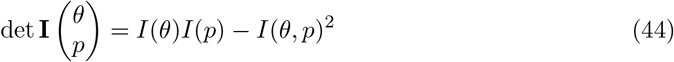

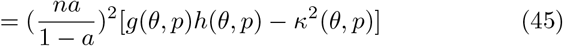

Finally, the variance of 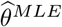 reads:

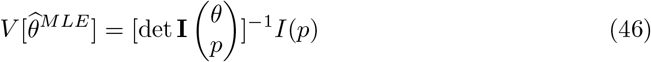

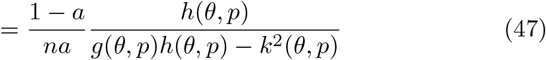

and its asymptotic variance:

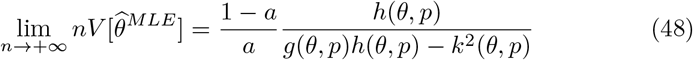

### 2.2 Detailed application of Proposition 1

Let us consider the following independent random variables

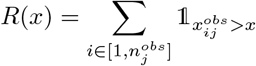

and

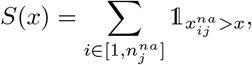

where 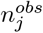 is the number of observed values in the sample *j*; and 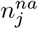 the number of missing values in this same sample.

Under Ass. 2 (the abundance values of peptides are independently distributed in the sample *j*), *R*(*x*) and *S*(*x*) are two independent binomial random variables:

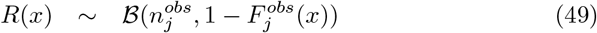

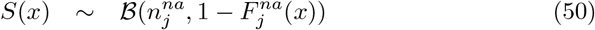

To match the notations of Prop. 1, we write:

- 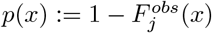
- 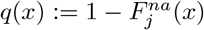
- 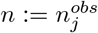
- 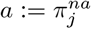 (i.e. 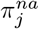 is the proportion of missing values in the sample *j*)

Therefore, the function *π*(*x*) defined in Eq. (8) of the article, which we recall reads as follow:

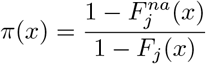

can be rewritten as

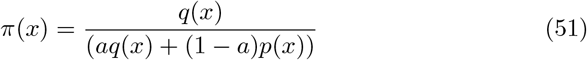

Moreover, 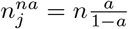. so that:

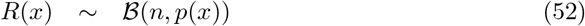

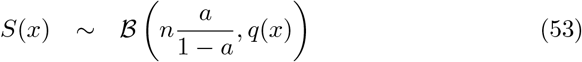

Thus, it is possible to apply Prop. 1 to define the maximum likelihood estimate of *π*(*x*).

### 2.3 Proof of Proposition 2

#### Proposition 2.

*Let*

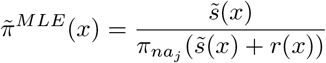

*where* 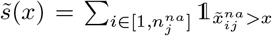 *and* 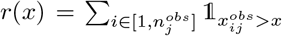. *Under Ass. 2 and Cor. 3, the proportion of missing values π_na_j__ is fixed. Then, for* 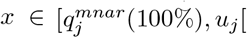,

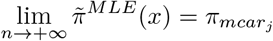

*Proof:* When

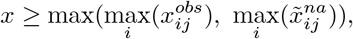

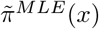 becomes undefined. Furthermore, when

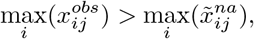

then 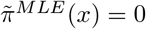 for 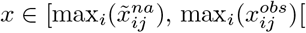. Conversely, when

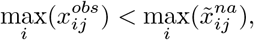

then 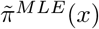 is undefined for 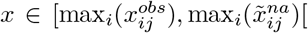. Thus, to ensure the consistency of the estimator, *x* must belong to 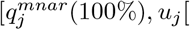, where

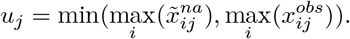

Therefore, for any 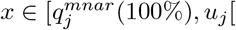, and when:

- 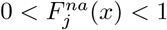
- 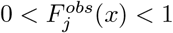
- the proportion of missing values *π_na__j_* is fixed,

we can derive from Cor. 3 and Prop. 1, that 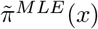 is an asymptotically unbiased estimate for *π_mcar_j__* when *n* → +∞.

### 2.4 Proof of Proposition 3

#### Proposition 3.

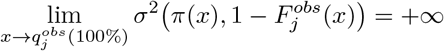

*Proof:* When *u* ≠ 0 and *υ* → 0, the functions *g, h* and *κ* (defined in Prop. 1) have the following asymptotic behavior:

- *g*(*u, υ*) → 0,
- *h*(*u, υ*) → +∞,
- *κ*(*u, υ*) → (1 − *π_na_j__*)/(1 − *π_na_j_ u_*)^2^.

Knowing that

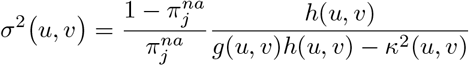

we derive that, 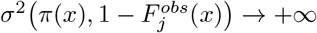 when 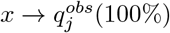

